# Uncoupling de novo pyrimidine biosynthesis from mitochondrial electron transport by ectopic expression of cytosolic DHODH

**DOI:** 10.1101/2024.08.09.607333

**Authors:** Andrea Curtabbi, Rocío Sanz Cortés, José Antonio Enríquez

## Abstract

Dihydroorotate dehydrogenase (DHODH) is an enzyme involved in the biosynthesis of pyrimidine nucleotides. In most eukaryotes, this enzyme is bound to the inner mitochondrial membrane, where it couples the synthesis of orotate with the reduction of ubiquinone. As ubiquinone must be regenerated by respiratory complex III, pyrimidine biosynthesis and cellular respiration are tightly coupled. Consequently, inhibition of respiration leads to cessation of DNA synthesis and impairs cell proliferation. We show that expression of *Saccharomyces cerevisiae* URA1 gene (*ScURA*) in mammalian cells uncouples biosynthesis of pyrimidines from mitochondrial electron transport. ScURA forms a homodimer in the cytosol that uses fumarate instead of ubiquinone as the electron acceptor, enabling oxygen-independent pyrimidine biosynthesis. Cells expressing *ScURA* are resistant to drugs that inhibit complex III and the mitochondrial ribosome. ScURA enables the growth of mtDNA-lacking ρ^0^ cells in uridine-deficient medium and ameliorates the phenotype of cellular models of mitochondrial diseases. This genetic tool uncovers the contribution of pyrimidine biosynthesis to the phenotypes arising from electron transport chain defects.

## Introduction

Cells obtain pyrimidine nucleotides for DNA and RNA synthesis either by de novo biosynthesis or by salvage of preformed bases ^1^. Dihydroorotate dehydrogenase (DHODH) catalyzes the fourth step of the biosynthetic pathway, oxidation of dihydroorotate to orotate. In most eukaryotes, DHODH is bound to the external surface of the inner mitochondrial membrane, where it uses ubiquinone (UQ) as an electron acceptor. Once UQ is reduced by DHODH, it diffuses into the internal mitochondrial membrane, where it is re-oxidized by respiratory complex III. Therefore, DHODH activity couples pyrimidine biosynthesis with the mitochondrial electron transport chain (mETC) ^2^. The biochemical consequence of this coupling is that de novo pyrimidine biosynthesis is not possible in absence of mitochondrial electron transport, due to the lack of UQ. This explains why cells lacking mtDNA or treated with mitochondrial inhibitors affecting complexes III or IV must rely solely on the salvage pathway, and they need exogenous uridine supplementation to grow ^3–5^.

Organisms capable to grow in absence of oxygen must produce orotate through a different mechanism. Here, DHODH is a cytosolic enzyme that uses either NAD^+^ or fumarate as electron acceptors. In these organisms, pyrimidine biosynthesis can thus proceed independently of cellular respiration. In yeast, the distribution of DHODH isoenzymes across various species appears to mirror their growth preferences: aerobic species harbor a mitochondrial DHODH, whereas facultative anaerobes possess a cytosolic DHODH ^6^. Anaerobic yeasts seem to have acquired the cytosolic DHODH gene by horizontal transfer from a bacterial source, and to subsequently have lost their eukaryotic gene ^7^. Oxygen-independent pyrimidine biosynthesis is therefore one of the main adaptations that enabled anaerobic growth ^8^.

The reason why evolution favored coupling of pyrimidine biosynthesis with the mitochondrial electron transport chain remains unknown. In this work, we asked whether it is possible to uncouple these two processes by expressing the gene encoding for cytosolic DHODH in mammalian cells.

## Results

In aerobic organisms, DHODH shuttles electrons via ubiquinone to complex III and functions as a respiratory enzyme (Figure 1A). Consequently, DHODH-mediated oxygen consumption is sensitive to inhibitors of both complex III (e.g. antimycin) and DHODH (e.g. brequinar). Other unicellular organisms have evolved different versions of DHODH, which utilize either fumarate or NAD^+^ as electron acceptors to oxidize dihydroorotate (DHO). These isoenzymes typically lack a mitochondrial import signal and a transmembrane domain, residing instead in the cytosol (Figure 1B).

**Figure 1.**
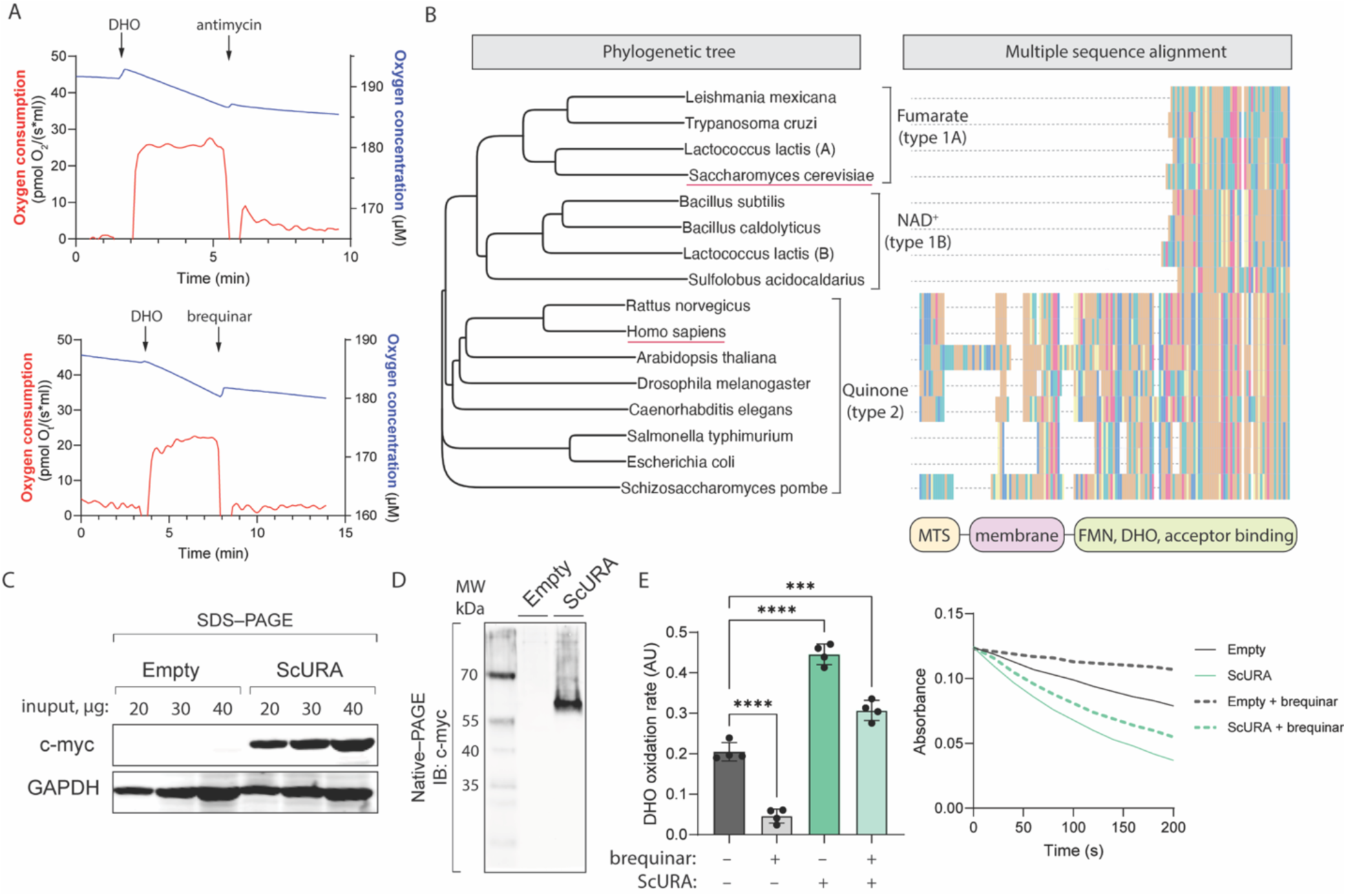
ScURA expression provides an alternative pathway for dihydroorotate oxidation. **(A)** Oxygen consumption of isolated mouse kidney mitochondria. Dihydroorotate (DHO) was given as a substrate and respiration was stopped by either complex III inhibition (upper panel) or DHODH inhibition (lower panel). **(B)** Phylogenetic tree and corresponding sequence alignment built from the aminoacidic sequences of DHODH isoforms from various phyla. The three branches reflect the differences between the electron acceptor used by DHODH and the type assigned by reference. *H. Sapiens* and *S. cerevisiae* are underscored in red. The functional domains are shown at the bottom, highlighting the lack of the mitochondrial targeting sequence (MTS) and of the transmembrane domain in type 1A and type 1B DHODHs. In the sequence alignment, only the first 200 amino acids are shown. **(C)** Immunoblot showing the expression of the myc-tagged ScURA protein in *ScURA* expressing 143B cells. **(D)** The same sample as in Figure 1C was run in native conditions, without SDS. The resulting immunoblot shows the myc-tagged ScURA dimer. (E) DHODH-dependent DHO oxidation by cell homogenates from ScURA expressing or control 143B cells. The DCPIP reduction rates per minute (left panel, see methods) and linear slopes of light absorbance at 600 nM from the first 200 seconds (right panel) are shown. Bars are mean ± SD. Ordinary one-way ANOVA, Tukey’s multiple comparison test, p<0.001 (***), p<0.0001 (****).

To enable mETC-independent de novo pyrimidine biosynthesis, we decided to ectopically express the URA1 gene from *S. cerevisiae* (*ScURA*) in mammalian cells. The yeast gene’s cDNA was codon-optimized for mammalian expression and a myc tag was added at the C-terminus through a flexible (GGS)3 linker to prevent steric interference between the tag and the enzyme. The gene was cloned in a lentiviral vector and an empty vector was used as the control for all subsequent experiments. In human 143B cells, *ScURA* expression yielded a product consistent with the predicted size of 32 kDa (Figure 1C). Previous studies have shown that URA1 protein forms homodimers in the cytosol of *S. cerevisiae* ^9^. Accordingly, when 143B homogenates were separated by native, non-denaturing, polyacrylamide gels electrophoresis, we observed that ScURA migrated at around 65 kDa, corresponding to the expected molecular weight of ScURA homodimer (Figure 1D). When we tested dihydroorotate dehydrogenase enzymatic activity in cell homogenates, we found that *ScURA* expressing cells possessed 2.25 times more activity than controls (Figure 1E). Approximately two third of this activity was insensitive to brequinar inhibition, as brequinar targets the ubiquinone binding site of human DHODH which is not present in ScURA ^10^. Therefore, ScURA expression provides an alternative pathway for DHO dehydrogenation in human cells.

Under normal cell culture conditions, DHODH activity is not a rate-limiting factor for cell proliferation and exogenous uridine supplementation is unnecessary (Figure 2A). Consistently, expression of *ScURA* did not affect cell proliferation or mitochondrial supercomplex organization in 143B cells (Figure 2B and Figure S1). To test whether *ScURA* could replace the endogenous DHODH activity in proliferating cells, we treated 143B cells with brequinar and we rescued pyrimidine biosynthesis inhibition by supplementing uridine in the growth medium. Brequinar suppressed cell proliferation in control cells, which was fully restored upon uridine supplementation (Figure 2C). On the other hand, in *ScURA* expressing cells, brequinar had no discernible effect on cell proliferation (Figure 2D). These results demonstrate that *ScURA* enables brequinar-insensitive de novo pyrimidine biosynthesis in human cells by bypassing mitochondrial DHODH.

**Figure 2.**
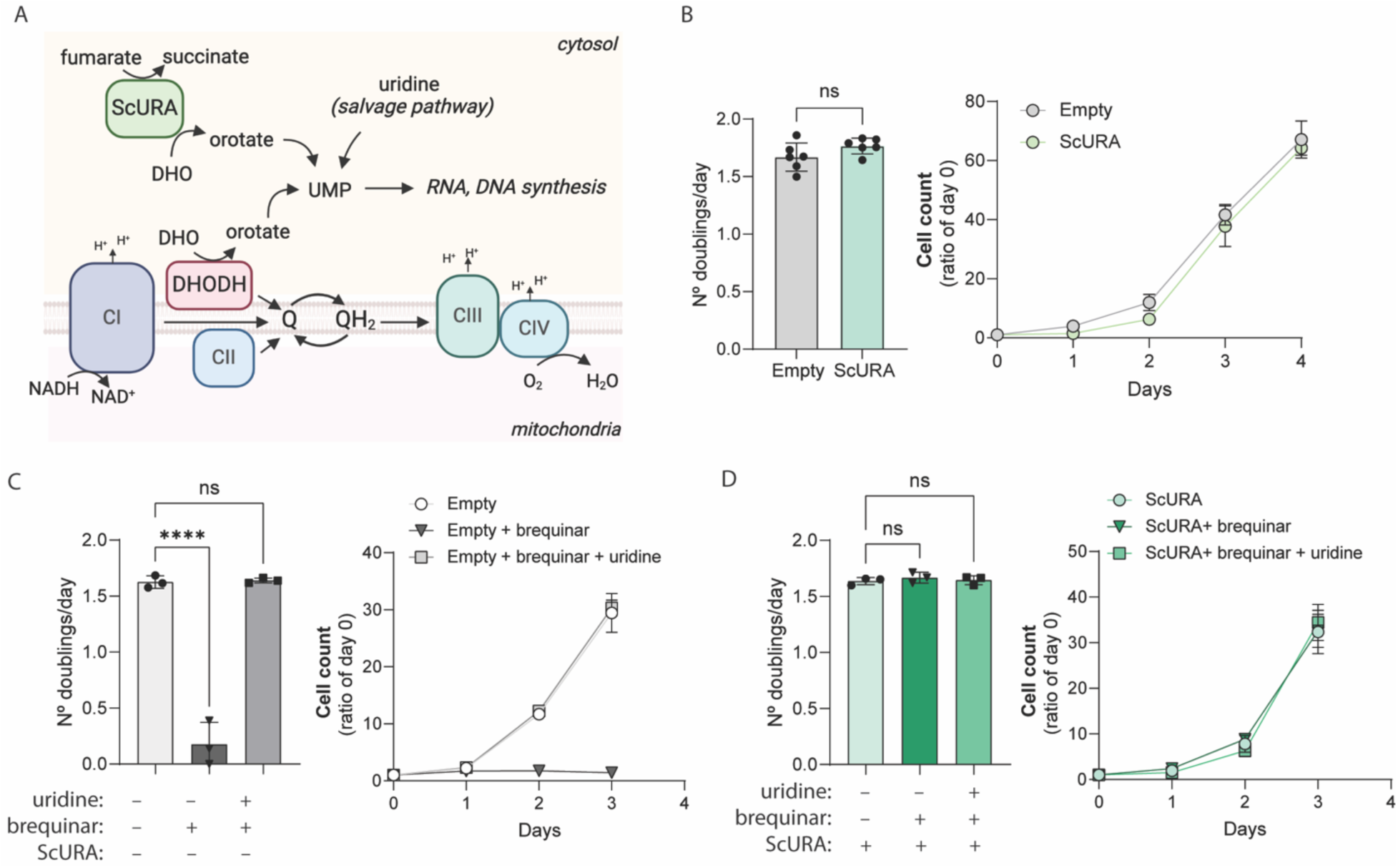
ScURA allows DHODH-independent pyrimidine biosynthesis. **(A)** Scheme showing the UMP production pathways by either mitochondrial DHODH, cytosolic ScURA or exogenous uridine. **(B)** Cell doublings (left) and corresponding growth curves (right) of ScURA expressing and control 143B cells grown in normal culture media. **(C)** Cell doublings (left) and corresponding growth curves (right) of control 143B cells treated with DHODH inhibitor brequinar and rescued by exogenous uridine supplementation in the growth media. **(D)** Same experiment as in Figure 2C, but with ScURA expressing 143B cells. Bars and dots are mean ± SD. Ordinary one-way ANOVA, Tukey’s multiple comparison test, p<0.0001 (****).

The dependency of pyrimidine biosynthesis on the mETC and was first discovered in chick embryo cells treated with chloramphenicol, which were found to be auxotrophic for uridine ^3^. Chloramphenicol, a mitochondrial ribosome inhibitor, impedes the assembly of respiratory complexes and the ATP synthase, blocking the electron flux through the mETC (Figure 3A). Therefore, we investigated whether ScURA expression could restore pyrimidine biosynthesis and rescue cell proliferation in cells treated with chloramphenicol. To this end, we cultured 143B cells in chloramphenicol and uridine for one week before assessing their ability to grow without uridine. While uridine supplementation was essential to maintain proliferation in control cells, *ScURA*-expressing cells were able to grow at the same rate in the absence of uridine (Figure 3B). We next sought to investigate whether ScURA could rescue cell proliferation when ubiquinol oxidation by complex III was disrupted. We treated cells with both antimycin A and myxothiazol, to completely block electron flow through complex III (Figure 3C) ^11^. Under these conditions, control cells were unable to proliferate without uridine, whereas the growth of *ScURA*-expressing cells was not dependent on uridine supplementation (Figure 3D). Therefore, ScURA allows mETC-independent de novo pyrimidine biosynthesis in human cells.

**Figure 3.**
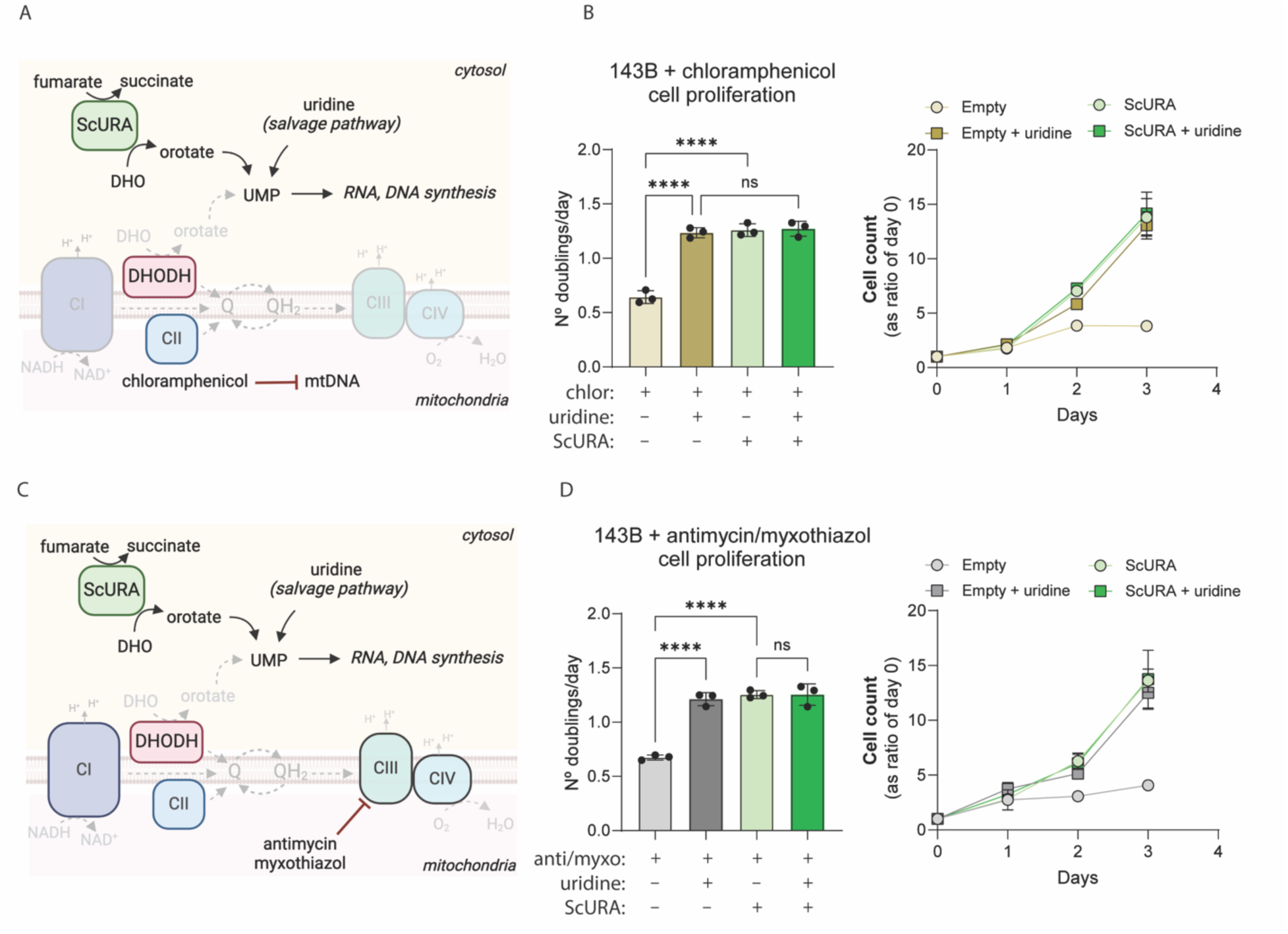
ScURA rescues cell proliferation upon mETC inhibition. **(A)** Scheme showing the UMP production pathways upon mitochondrial ribosome inhibition by chloramphenicol. **(B)** Cell doublings (left) and corresponding growth curves (right) of ScURA expressing and control 143B cells treated with chloramphenicol (chlor) and rescued with exogenous uridine supplementation. Cells were pre-treated with chloramphenicol for one week in the presence of uridine to devoid them of respiratory complexes (*13*). **(C)** Scheme showing the UMP production pathways upon respiratory complex III inhibition by antimycin A and myxothiazol. **(D)** Cell doublings (left) and corresponding growth curves (right) of ScURA expressing and control 143B cells treated with antimycin A and myxothiazol (anti/myxo) and rescued by exogenous uridine supplementation. Bars and dots are mean ± SD. Ordinary one-way ANOVA, Tukey’s multiple comparison test, p<0.0001 (****).

Cells depleted of mtDNA, known as ρ^0^ cells, are dependent on uridine and pyruvate for growth. We therefore asked whether we could revert uridine auxotrophy in ρ^0^ cells with ScURA. Transduction of two ρ^0^ cell lines, mouse L929 ρ^0^and human 143B ρ^0^, with the *ScURA* gene allowed growth in the absence of uridine, while maintaining their pyruvate dependence (Figure 4A and 4B). Together, these results show that de novo pyrimidine biosynthesis can be fully restored in cells suffering from various degree of mitochondrial dysfunction by *ScURA* expression.

**Figure 4.**
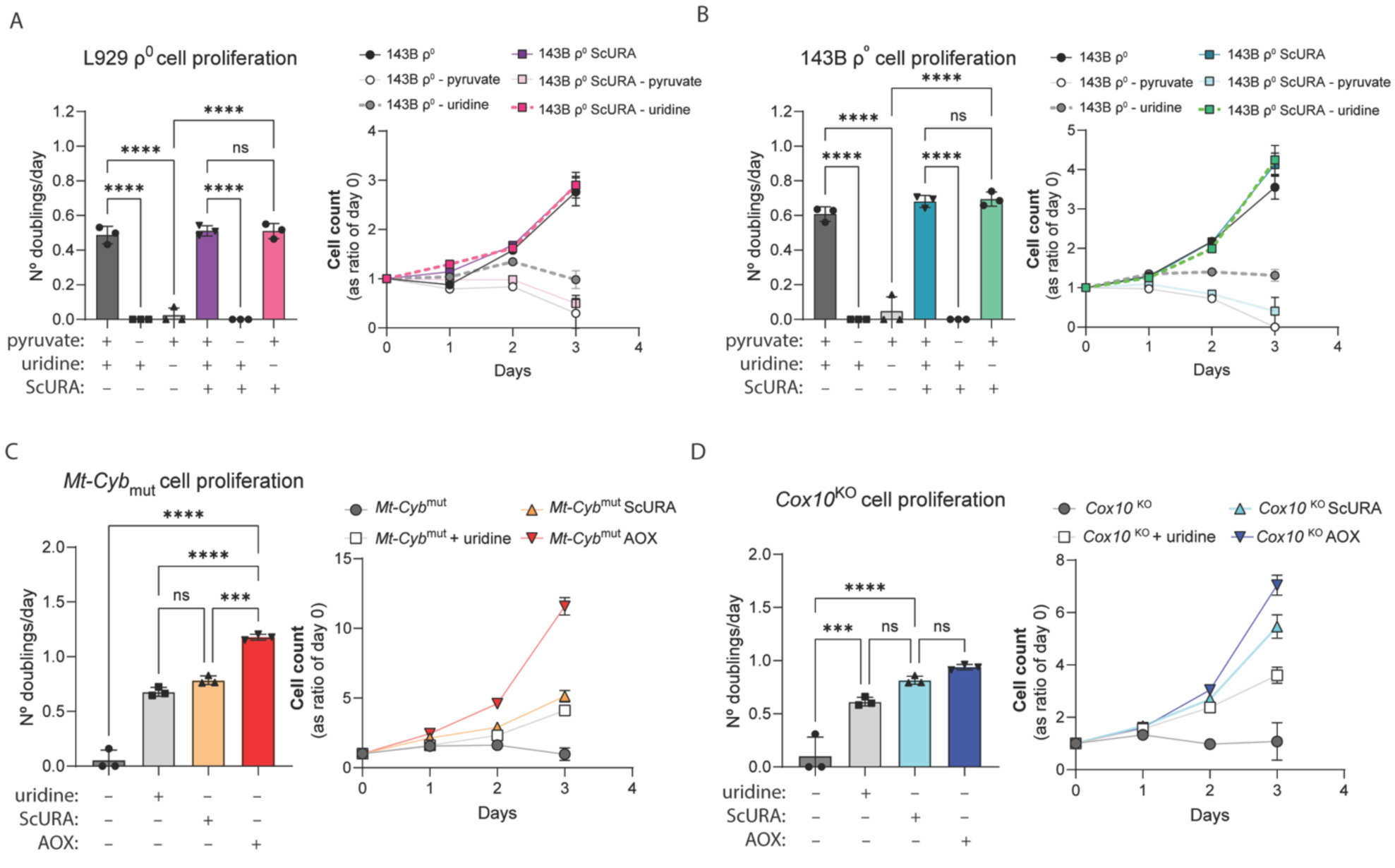
ScURA alleviates cell proliferation defects caused by mitochondrial dysfunction in mitochondrial disease models. **(A)** Cell doublings (left) and corresponding growth curves (right) of ScURA-expressing and control mouse L929 p^0^ cells. Exogenous uridine and/or pyruvate was added to the growth media where indicated. **(B)** Same experiment as Figure 4A but with ScURA-expressing and control human 143B p^0^ cells. **(C)** Cell doublings (left) and corresponding growth curves (right) of ScURA expressing, AOX expressing and control *Mt-Cyb*^mut^ cells. (D) Same experiment as Figure 4A but with ScURA-expressing, AOX-expressing and control *Cox10*^KO^ cells. Bars are mean ± SD. Ordinary one-way ANOVA, Tukey’s multiple comparison test, p<0.001 (***), p<0.0001 (****).

Mitochondrial diseases are a heterogeneous group of maladies with extremely variable manifestations, are poorly predictable and often lack an effective treatment. Cell lines derived from patients suffering from defects in respiratory complexes benefit from uridine supplementation ^12^. Expression of the alternative oxidase (AOX) from *C. intestinalis* in human 293T confers cyanide resistance to cells ^13^. Equally, expression of AOX from *E. nidulans* in mouse L929 ρ^0^ cells rescues uridine auxotrophy ^14^. In mouse and human cells deficient for CIII or CIV, AOX can also rescue uridine auxotrophy, ameliorating their metabolic phenotype ^15, 16^. In these scenarios, AOX restores the electron flux through the mETC bypassing respiratory complex III and complex IV, allowing pyrimidine biosynthesis ^14^. To examine the implications of *ScURA* expression in cellular models of mitochondrial diseases we introduced it in *Mt-Cyb*^mut^ cells, which lack respiratory complex III due to a point mutation in the mtDNA *cyb* gene ^17^, and in *Cox10*^KO^ cells, which cannot assembly respiratory complex IV due to the absence of the COX10 assembly factor ^18^ (Figure S2). *ScURA* expression enabled the growth of both cell lines in the absence of uridine (Figure 4C and 4D). We concluded that the growth defect caused by pathological mutations in mitochondrial genes can be alleviated by ScURA.

## Discussion

The ectopic expression of enzyme orthologs from other phyla has been a valuable tool for the investigation of the mitochondrial electron transport chain. For example, *Lactobacillus brevis* NADH oxidase generates NAD^+^ bypassing complex I, and has been used extensively both in cellular and animal models ^19, 20^. *C. intestinalis AOX*, which substitute complex III and IV ^13^, and *S. cerevisiae NDI1*, which substitute complex I ^21^ are being explored as therapeutic approaches for mitochondrial diseases ^3, 22^. We have extended this genetic toolkit with ScURA, which provides a mETC-independent pathway for orotate production.

The reason why most eukaryotes rely on a mitochondrial DHODH is unknown. The connection between DHODH and mitochondrial electron transport could help to fine tune key metabolic processes such as DNA and RNA synthesis with oxygen and nutrient availability. When oxygen supply is limited, reduction in complex III and IV activity would deplete the UQ pool, slowing down pyrimidine biosynthesis and cell growth ^23^. Decreased pyrimidines level might, in turn, stimulate respiratory supercomplexes assembly by ether phospholipid accumulation ^24^. A mitochondrial DHODH might therefore support proliferation only in favorable environments.

It has been shown that during hypoxia ubiquinol can be oxidized by the reversed activity of succinate dehydrogenase, allowing complex-III independent DHODH catalysis ^25^. However, the extent of this pathway’s contribution to pyrimidine biosynthesis appears to be minimal when complex III activity is inhibited either genetically or pharmacologically *in vitro.* Under such conditions, cell proliferation is almost completely halted in absence of uridine supplementation or ScURA expression. Further investigation is needed to fully understand the function of this metabolic coupling.

ScURA allowed us to evaluate contribution of de novo pyrimidine biosynthesis to cellular growth independently from cellular respiration. In recent years, genetic-engineered mouse models have enabled the study of a wide spectrum of cell type-specific functions. With this approach, the integrity of the mitochondrial electron transport chain has been shown to be essential for hematopoiesis ^26^, angiogenesis ^27^, regulatory and NK T cells functions ^28^ and many more. Such dependence is often assayed by genetic disruption and pharmacologic inhibition of single mitochondrial respiratory complexes. By this approach, the activity of all UQ-dependent dehydrogenases is blocked, raising the question of which mETC-dependent process is the responsible for the observed phenotype. The relative contribution of each UQ-dependent pathway can be assessed only by ectopic expression of UQ-independent dehydrogenases *in vivo* ^19, 27^. In this way, the development of effective and precise pharmacological intervention strategies can be undertaken following a precise mechanistic rationale. We designed ScURA following this same logic.

One key issue which we think ScURA will help to disentangle is on the metabolic requirements of tumor growth. Tumor cells in which the integrity of mETC is disrupted, cannot form tumors when injected subcutaneously ^29, 30^. Pyrimidine biosynthesis has been suggested to be the main limiting factor of respiratory deficient tumor cells ^31^. ScURA expression in respiratory deficient tumor cells will provide a straightforward model to directly test this and other hypotheses *in vivo*.

## Author contributions

Conceptualization: AC & JAE. Data collection and analysis: AC and RSC. writing: AC & JAE.

## Acknowledgments

We acknowledge all GENOXPHOS group members for their scientific discussions contributing to this manuscript. We thank members of the CNIC facilities (Cell culture and Viral vectors) for their technical assistance.

## Material and Methods

### Cell culture

Cells were grown in complete DMEM medium (D5796, containing 4500 mg/L glucose, 2 mM l-glutamine) supplemented with dialyzed 10 % fetal bovine serum (FBS, Sigma F7524), 1 % penicillin-streptomycin (PenStrep, Lonza), at 37 °C in an atmosphere of 5 % CO_2_/95 % air. Where indicated, 50 μg/mL uridine and/or 1 mM sodium pyruvate (Sigma) were added to the growing media. Inhibitors were purchased from Sigma and used as follows: brequinar (2 μM); antimycin A/myxothiazol (20 μM each); chloramphenicol (40 μg/mL). Control cells were treated with the appropriate vehicle (ethanol or DMSO). Origin of cell lines: ρ^0^ cells ^32^ *Mt-Cyb*^mut^ and in *Cox10*^KO^ AOX-expressing cells ^16^.

### Generation of ScURA expressing cells

The coding sequence of S. cerevisiae URA1 gene was obtained from ^33^ and modified as follows. A myc tag was added at the C-terminus through a (GGS)3 linker and at both ends restriction enzyme sites were inserted. Codon-optimization and oligonucleotide synthesis were performed by Genewiz. The gene was cloned in the pWXLd lentiviral vector (modified from Trono’s lab with a puromycin resistance gene, Addgene). Lentivirus production was carried out by the Viral Vector unit at CNIC. Viral supernatants were filtered and added to cells with 8 μg/ml polybrene overnight. Two days after transduction, ScURA expressing cells were selected with puromycin 0.5 μg/ml approximately for week, until non-transduced cells in a control plate were all dead. For each cell line, a cell line transduced with an empty vector was used as a control.

### Growth curve and cell doubling time

On day zero, 40,000 cells were plated in 6 well plates. For each condition, three wells were incubated 6 hours (to let cells adhere to the plate), collected, and counted. This count was recorded as the number of cells at day 0. For 3-4 days, fresh media and the appropriate treatments were added to each well every day. On the last day, three wells per condition were collected and counted. The number of doublings/day was calculated as = 24/ ([ T × (ln2 ) ] / [ ln (Nf / N0) ]), where T= time in hours, Nf= number of cells on the final day, N0= number of cells at day zero.

### Whole-cell homogenate and mitochondria isolation

Cells from 2 to 10 150 mm plates were collected with trypsin, washed in PBS and resuspended in cold sucrose buffer (0.32 M sucrose, 1 mM EDTA, 10 mM Tris-HCl, pH 7.4). Cells were then homogenized in a Teflon potter-type tissue homogenizer with 20-40 “pops”. The homogenate was centrifuged at 1000 g, 5 min, the nuclear pellet was discarded and the post nuclear whole-cell homogenate was stored at -80 °C. When mitochondria were isolated, the whole-cell homogenate obtained was further centrifuged at 10,000 g, 10 min. The pellet obtained was resuspended in sucrose buffer and stored at −80 °C. Mitochondria isolation from kidney of CD1 mice was carried out in the same way. In case mitochondria were used fresh for functional assays (ex. respirometry), sucrose buffer supplemented with bovine serum albumin 0.1 % was used throughout all homogenization steps.

### Spectrophotometric DHODH activity

DHODH activity was measured following the method of Knecht et al. ^34^ with some modifications. 100µg of whole-cell homogenates were used for each reaction, in final volume of 200 µl. Each biological replicate was split into three technical replicates. DHODH-dependent reduction of 2,6-dichloroindophenol (DCPIP) was measured with a UV–visible spectrophotometer equipped with a 96-well plate reader. Samples were resuspended in reaction buffer (Tris-HCl 50 mM, KCl 150 mM, KCN 3 mM, DCPIP 0.05 mM, decylubiquinone 0.25 mM, pH=7.5) and incubated for 2 minutes at 37°C. The reaction was started by addiction of dihydroorotate (DHO) 1mM. The first 3 minutes of the reaction were linear and were used for the analysis. The rate of background (DHODH-independent) DCPIP reduction was measured in parallel by omitting DHO from the reaction mixture and was subtracted from the DHO-dependent rate for each sample. Brequinar was added were indicated at 0.05 mM, controls were treated with the same amount of DMSO.

### Gel electrophoresis

All gels were prepared in house with Tris-HCl buffer and acrylamide/bis. Protein samples were quantified and loaded on the gel, and electrophoresis was developed in Tris-glycine solution. For denaturing electrophoresis, 0.1% SDS was added to the gels and protein samples were incubated for 1 min at 95 °C in loading buffer (Tris-HCl 50 mM pH 6.8, 2 % SDS, 10 % glycerol, 1 % β-mercaptoethanol, 0.02 % bromophenol blue) before loading. For non-denaturing native gels, SDS was not added to the gel, whole-cell homogenates were not boiled, and the loading buffer used did not contained SDS nor β-mercaptoethanol.

### Immunoblotting

Proteins were transferred to PVDF membrane (Immobilon-FL, 0.45 μm) by transfer in Bio Rad Mini Trans-Blot Cell or Trans-Blot Cell systems, in 48 mM Tris, 39 mM glycine, 20 % methanol transfer solution overnight at 30 V. The membrane was then blocked in 0.1 % PBS-tween, 5 % BSA solution for 1 h at room temperature and incubated with primary antibody overnight at 4 °C. After that, it was incubated with secondary antibody for 1 hour at room temperature. Membrane revealing was performed by using fluorescent secondary antibody and revealed with the Odyssey imaging system (LI-COR biosciences).

### BN-PAGE and Complex I In-gel activity

The procedure was carried on as detailed in ^35^. Briefly, 100 μg of mitochondria were incubated with 400 μg of digitonin for 5 min in 50 mM NaCl buffer, 50 mM imidazole, 5 mM aminocaproic acid at a final concentration of 10 μg/μl. Electrophoresis was performed in cold chamber and developed for half hour at 90 V with cathode buffer A. Then, the cathode buffer was exchanged for cathode buffer B and electrophoresis continued for approximately one more hour at 300 V. Measurement of NADH dehydrogenase activity of complex I was determined on the same gel after BN-PAGE electrophoresis by incubating the gel in 0.1 M Tris-HCl, pH 7.4, 0.14 mM NADH and 1 mg/ml NitroBlue tetrazolium solution at room temperature.

### Respirometry

Oxygen consumption was assayed using a Clark type polarographic oxygen sensor (Oroboros instruments) in a 2 ml isolated chamber, in agitation with a magnetic stirrer, at 37 °C, in Mir 05 respiration medium (Oroboros instruments). 0.5 mg of kidney mitochondria were incubated with the indicated substrates and inhibitors.

### Phylogenetic tree construction

The aminoacidic sequence of DHODH from various species was retrieved from uniport. The R packages msa ^36^, ape ^37^ and ggmsa ^38^ were used for multiple sequence alignment, phylogenetic tree construction and generation.

---

**Figure S1.**
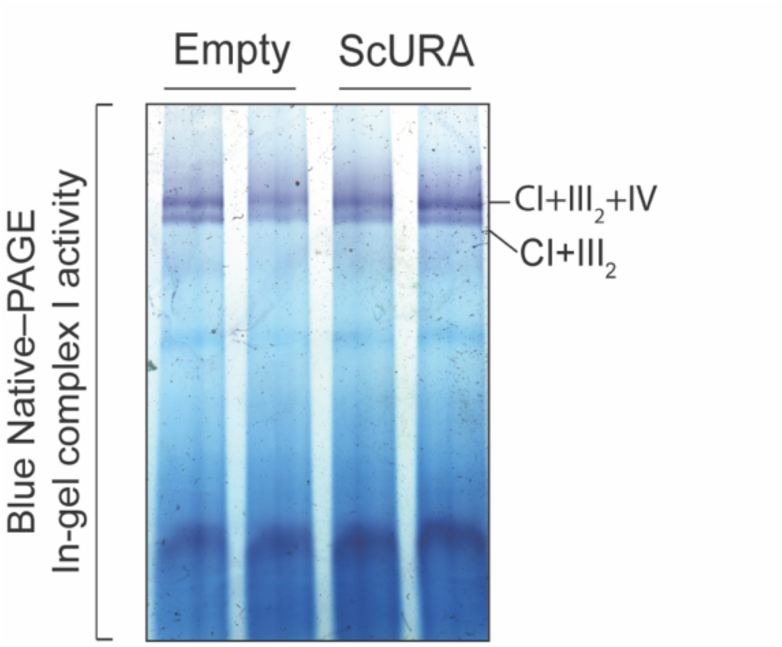
In-gel complex I activity of mitochondria isolated from ScURA expressing and control 143B cells. Related to Figure 2.

**Figure S2.**
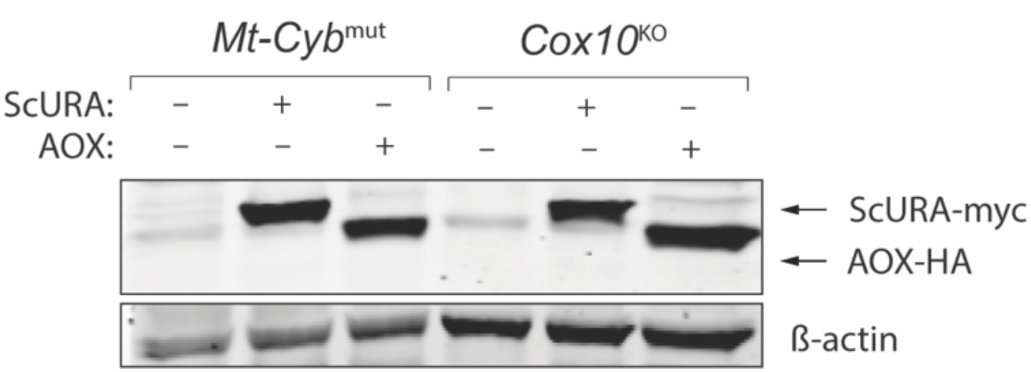
Immunoblot showing c-myc tagged ScURA and HA-tagged AOX expression in *Mt-Cyb*^mut^ and *Cox10*^KO^ cells. Related to Figure 4.

